# Epigenetic deregulation of lamina-associated domains in Hutchinson-Gilford Progeria Syndrome

**DOI:** 10.1101/520403

**Authors:** Florian Köhler, Felix Bormann, Günter Raddatz, Julian Gutekunst, Tanja Musch, Frank Lyko, Manuel Rodríguez-Paredes

## Abstract

Hutchinson-Gilford Progeria Syndrome (HGPS) is a progeroid disease characterized by the early onset of some classically age-related phenotypes including arthritis, loss of body fat and hair and atherosclerosis. Cells from affected individuals express a mutant version of the nuclear envelope protein Lamin A (termed Progerin) and have previously been shown to exhibit prominent chromatin changes. Here, we identify epigenetic deregulation of lamina-associated domains (LADs) as a central feature in the molecular pathology of HGPS. Using ATAC-see/-seq and Infinium MethylationEPIC BeadChip-mediated DNA methylation profiling, we demonstrate that dermal fibroblasts from HGPS patients exhibit both chromatin accessibility and DNA methylation changes that are enriched in LADs. Importantly, we further show that these epigenetic alterations are associated with HGPS-specific gene expression changes. Together, our results establish a central involvement of LADs in the epigenetic deregulation of HGPS and provide novel insight into the molecular changes associated with the disease.

## Introduction

The nuclear lamina is a filamentous mesh network that lines the inner nuclear membrane of metazoan nuclei. Its main component are type V intermediate filaments termed lamins, four of which, Lamin A, B1, B2 and C, are expressed in metazoan cells^1,2^. Over 180 mutations causing at least 13 different diseases, collectively called laminopathies, have been described for their corresponding genes^3^. One of the most severe laminopathies is Hutchinson-Gilford Progeria Syndrome (HGPS), a rare progeroid disease characterized by the early onset of some classically age-related phenotypes including osteoporosis, alopecia, loss of body fat and hair, and atherosclerosis^4,5^. Affected individuals are usually diagnosed within the first year of life due to a failure to thrive, and suffer from rapid disease progression with death occurring in their teens as a consequence of myocardial infarction or stroke^5,6^.

The classical form of HGPS is caused by an autosomal dominant mutation in exon 11 of the *LMNA* gene, which encodes both Lamin A and C^7^. The mutation activates a cryptic splice site, causing the expression of a mutant Lamin A that lacks 50 amino acids near the C-terminus while leaving Lamin C unaffected^7,8^. The resulting truncated protein, referred to as Progerin, undergoes aberrant posttranslational modification and retains a farnesyl residue at its C-terminal CaaX motif, thus becoming permanently associated with the nuclear lamina and causing characteristic morphological changes^6^. Progerin-expressing cells have been shown to display a wide range of cellular defects such as premature cellular senescence, increased levels of reactive oxygen species, clustering of nuclear pores, delayed DNA repair and shortened telomeres^6,9–14^. Interestingly, low level Progerin expression has also been found in cells from normal, aged individuals, suggesting that it might play a role in normal aging as well^15–17^.

HGPS cells also exhibit a number of epigenetic aberrations. Most prominently, heterochromatin markers such as the histone modifications H3K27me3 and H3K9me3, as well as heterochromatin protein 1 (HP1) and the H3K27me3 methyltransferase EZH2, have been shown to be downregulated in fibroblasts from HGPS patients^15,18,19^. Conversely, H4K20me3, another heterochromatin mark, has been reported to be increased in HGPS cells and after ectopic Progerin expression^18,20^. Furthermore, Hi-C experiments indicate that late-passage HGPS cells lose the spatial compartmentalization of active and inactive chromatin domains present in healthy cells^19,21^.

Less is known about the HGPS-associated role of another key epigenetic modification, DNA methylation. An earlier comparison of BJ and HGPS skin fibroblasts using bisulfite padlock probes, a method for the targeted quantification of DNA methylation at a limited number of CpGs^22^, identified 586 differentially methylated autosomal genes in HGPS^23^. Another study reported profound DNA methylation changes in a set of age-related genes in HGPS patients^24^. However, the authors used samples of adult onset, i.e., non-classical progeria, for their comparisons^24^, thus leaving the question of DNA methylation alterations in classical HGPS unanswered. Finally, it was recently demonstrated that HGPS fibroblasts from some patients exhibit an increased ‘DNA methylation age’ (an estimate of the biological age computed on the basis of the methylation status of 391 genomic loci)^25^, hence suggesting a considerable degree of underlying epigenetic changes. A comprehensive characterization of these aberrations in classical HGPS, however, remains to be performed.

A potential candidate for epigenetic changes in HGPS are lamina-associated domains (LADs), i.e., regions of the DNA that are in close contact with the nuclear lamina. They are considered to assist in the spatial organization of the genome and to exert a role in gene repression^26–28^. Importantly, LADs largely overlap with late-replicating regions^26,27^, are mostly gene-poor^26^ and are enriched for the heterochromatin marks H3K9me2 and H3K9me3^26,29^. Despite their heterochromatic nature, LADs are not associated with high levels of cytosine methylation. In fact, they overlap to a large extent with partially methylated domains (PMDs)^30,31^, i.e., long stretches of DNA with reduced levels of DNA methylation. Interestingly, PMDs have also been shown to undergo hypomethylation as a consequence of mitotic cell division and cell aging^32^. Whether they also become differentially methylated in HGPS is still an open question.

Here, we identify epigenetic deregulation of LADs as a central feature of the epigenetic alterations in HGPS. Using ATAC-see/-seq and Infinium MethylationEPIC BeadChip-mediated DNA methylation profiling, we demonstrate that dermal fibroblasts from HGPS patients exhibit both chromatin accessibility and DNA methylation changes that are enriched at LADs. Importantly, we further show that these epigenetic alterations are associated with HGPS-specific gene expression changes. Together, our results strongly implicate epigenetic deregulation of LADs as an important and previously unrecognized key feature of HGPS.

## Results

### ATAC-see and ATAC-seq reveal genome-wide chromatin accessibility changes enriched in lamina-associated domains (LADs)

We obtained primary skin fibroblasts from nine progeria patients and six controls (Suppl. Tab. 1). We tested all these cell lines for characteristic nuclear morphology changes and Progerin expression at the transcript and protein levels. Specifically, HGPS fibroblasts exhibited a wide range of nuclear malformations including characteristic wrinkling and lobulation of the nuclear lamina (Suppl. Fig. 1A). Importantly, the fraction of cells showing nuclear malformations ranged from 30-89% in HGPS cells, but only from 3-15% in control cells (Suppl. Fig. 1B). In addition, expression of Progerin as a consequence of an upregulation of the Δ150 *LMNA* transcript was readily detected in all HGPS, but not in control samples (Suppl. Fig. 1C-F).

In order to obtain further insight into the extent of epigenetic aberrations in HGPS cells, we imaged the transposase-accessible genome in HGPS and control fibroblasts using ATAC-see^33^. Through the insertion of fluorophores by the Tn5 transposase at open chromatin sites, this technique allows the visualization of accessible chromatin using microscopy. Interestingly, whereas bright foci representing highly accessible chromatin regions were detected in 82-91% of control cell nuclei, only 34-53% of HGPS nuclei revealed these foci (Figs. 1A and 1B, P=1.43e-2, unpaired t-test). Moreover, the number of foci was negatively correlated with the number of malformed nuclei in HGPS cells (R^2^= 0.30), suggesting that HGPS-related nuclear malformation has an effect on chromatin accessibility (Fig. 1C).

**Figure 1.**
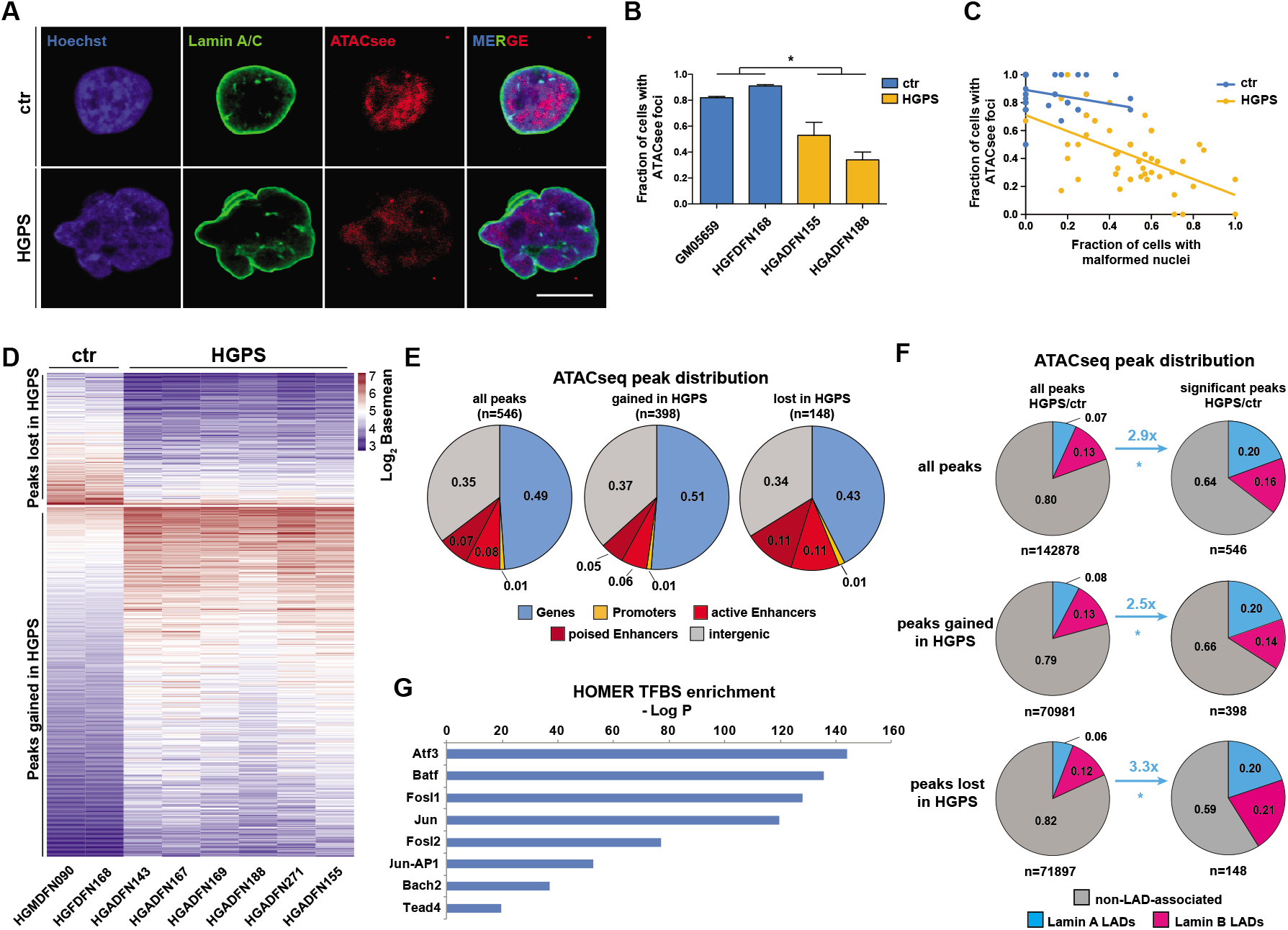
ATAC-see and ATAC-seq reveal genome-wide chromatin accessibility changes enriched in lamina-associated domains (LADs). **(A)** ATAC-see reveals loss of highly accessible chromatin foci in malformed HGPS nuclei. Scale bar = 10 μm. **(B)** Quantification of cells with 3 or more ATAC-see foci (P=1.43e-2, unpaired t-test). 50 nuclei were counted per sample (three replicates). **(C)** Correlation of the number of cells with 3 or more ATAC-see foci with the number of cells with malformed nuclei (HGPS: R^2^=0.30). Malformed nuclei were quantified as in (B). **(D)** Regions gaining (n=398) or losing (n=148) accessibility in HGPS compared with controls (q<0.05, Benjamini-Hochberg). **(E)** Distribution of ATAC-seq peaks across genes, promoters and enhancers. **(F)** Distribution of ATAC-seq peaks across Lamin A-, Lamin B-and non-LAD-associated regions (*P<0.05, Fisher’s Exact test). **(G)** HOMER transcription factor motif enrichment analysis reveals members of the AP1 family are highly enriched (q<0.01, Benjamini-Hochberg) in differentially accessible regions.

To further investigate this possibility and to quantify the magnitude of changes in HGPS fibroblasts, we globally profiled the accessible genome in HGPS cells using ATAC-seq. This identified 546 significantly differentially accessible regions between HGPS and control fibroblasts, of which 398 and 148 gained and lost accessibility in HGPS, respectively (q<0.05, Benjamini-Hochberg, Fig. 1D and Suppl. Fig. 2A). About one half of the differentially accessible regions were located in genes, about one third mapped to intergenic regions and a minor fraction to active and poised enhancers (Fig. 1E).

Interestingly, we found several lines of evidence suggesting that genomic regions in contact with the nuclear lamina, especially those associated with Lamin A, undergo substantial alterations in chromatin accessibility in HGPS cells. First, Lamin A-, but not Lamin B-associated LADs were enriched 2.5- (P=5.62e-15, Fisher’s Exact test) and 3.3-fold (P<9.44e-09, Fisher’s Exact test) among the regions gained and lost in HGPS, respectively (Fig. 1F). Second, regions associated with the active chromatin marks H3K4me3 (P<2.20e-16), H3K27ac (P=2.30e-3) and H3K36me3 (P=5.88e-06) were significantly underrepresented among the differentially accessible regions, whereas regions associated with the repressive chromatin mark H3K9me3 (P=3.70e-4) were slightly overrepresented (Fisher’s Exact test for all, Suppl. Fig. 2B). Third, we observed that the relatively gene-poor chromosome 18, which tends to be located near the nuclear periphery and shows multiple LAD contacts in proliferating cells^34,35^, exhibited more than 7 times more differentially accessible regions than the similarly sized, gene-rich, and more centrally located chromosome 19 (Suppl. Fig. 2C). Finally, we found that binding sites of members of the Activator Protein 1 (AP1) family of transcription factors, which have previously been shown to be associated with the nuclear lamina in mammalian cells^36^, are highly enriched in the differentially accessible regions (q<0.01, Benjamini-Hochberg, Fig. 1G). As Fos/Jun transcription factors regulate a wide range of cellular processes including cell proliferation, cellular differentiation and apoptosis^37,38^, this is likely to have functional implications in HGPS cells. Taken together, our ATAC-see and ATAC-seq experiments provide evidence that the chromatin accessibility landscape is substantially altered in HGPS cells, with Lamin A-associated LADs being at the center of these changes.

### DNA methylation profiling in HGPS reveals two patient subgroups and LAD-enriched hypermethylation

Loss of DNA methylation in lamina-associated, late-replicating regions known as PMDs has recently been shown to constitute a biomarker of cellular aging^32^. To investigate whether the LAD-specific chromatin accessibility changes in HGPS cells are accompanied by alterations in DNA methylation, we performed DNA methylation profiling using Infinium MethylationEPIC BeadChips, which capture the methylation status of more than 850,000 CpGs in the human genome.

After batch correction and removal of sex chromosome-associated probes we identified 19,759 differentially methylated (P<0.05, F-test) probes between nine HGPS and six control samples (Fig. 2A). Unsupervised consensus clustering of the 5,000 most variably methylated probe clusters separated the samples into three groups: while the controls formed one uniform group, the HGPS samples were split into two subgroups (Fig. 2B and Suppl. Fig. 3A). This substructure was not associated with patient age, body site of sampling, sex, strength of Progerin expression or passage number. However, it was in agreement with a subclassification of HGPS samples based on a recently reported DNA methylation age acceleration in some HGPS patients^25^. When we applied the corresponding age estimator to our sample set, the samples were divided into an age-accelerated group (median Δ age = 9.73 years) and a group with a slight age deceleration (median Δ age = −1.51 years), which overall matched the two subgroups identified through consensus clustering (Fig. 2C). This finding suggests that individual HGPS patients differ significantly with regard to their DNA methylation patterns and that this epigenetic modification may represent a novel tool for the classification of affected individuals.

**Figure 2.**
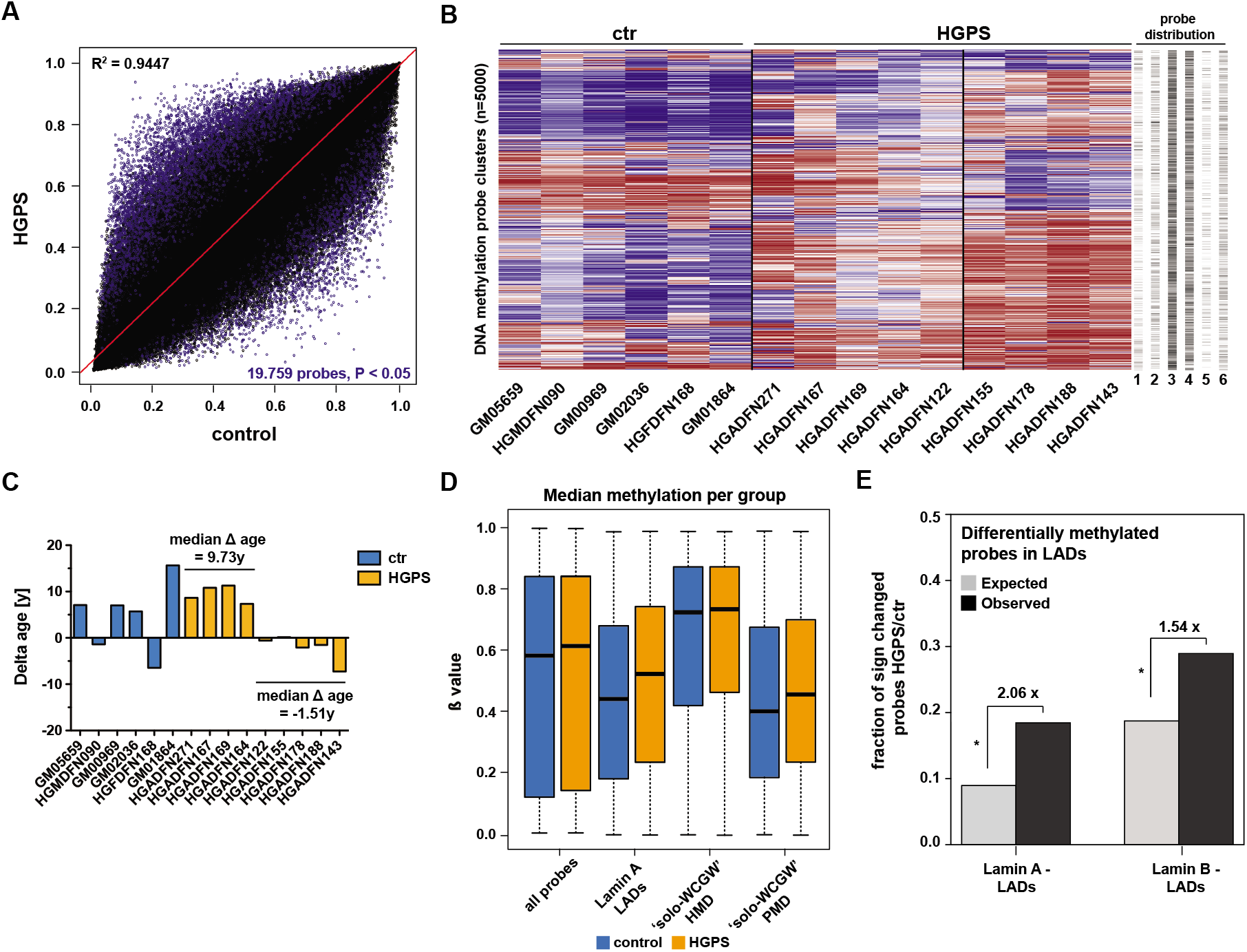
DNA methylation profiling in HGPS reveals two patient subgroups and lamina-associated domain (LAD)-enriched hypermethylation. **(A)** Scatter plot comparing the methylomes of HGPS and control fibroblasts. Differentially (P < 0.05, F-test) methylated probes are shown in blue. **(B)** Consensus clustering based on 5000 most variable probe clusters between HGPS and control samples (1=CpG islands, 2=promoter, 3=gene body, 4=intergenic, 5=Lamin A-assoc., 6=Lamin B-assoc.). β values are colored from blue (β = 0) to red (β = 1). **(C)** Difference (= Δ age) between DNA methylation age (Skin&Blood Clock^25^) and chronological age for all samples. **(D)** Differential (β value) methylation of Lamin A LAD-(P<2.20e-16), soloWCGW HMD-(P=4.42e-15) and soloWCGW PMD-(P<2.20e-16) associated probes (all: Welch Two Sample t-test) with median indicated as a black line. HMD=Highly Methylated Domain, PMD=Partially Methylated Domain. **(E)** Enrichment of LAD-associated probes among the probes differentially (P<0.05, F test) methylated between HGPS and control samples (Lamin A: 2.06-fold, Lamin B: 1.54-fold, both: P<2.20e-16, Fisher’s Exact test). Expected numbers were calculated based on the fraction of LAD-associated probes among all probes normalized to the number of differentially methylated probes.

Of note, most of the 5,000 most variable probe clusters used for the subgrouping contained probes located in genes and intergenic regions, while only a few were located in CpG islands and promoters (Fig. 2B). Together with the fact that CpG island methylation in the HGPS samples was virtually unchanged (Suppl. Fig. 3B), this indicates that aberrant CpG island methylation, a prominent epigenetic alteration of normal cells during aging^39,40^, plays only a minor role in HGPS.

One of the major epigenetic alterations known in HGPS is the loss of heterochromatin-associated histone marks at the nuclear periphery of Progerin-expressing cells^15,18,19^. We therefore tested whether DNA methylation is similarly altered in genomic regions associated with the nuclear lamina. While we only found a small increase in median methylation levels of all 850,000 EPIC probes, Lamin A LAD-associated probes exhibited a strong and significant (P<2.20e-16, Welch Two Sample t-test) increase (Fig. 2D) and were enriched 2.06-fold among the differentially methylated probes in HGPS (P<2.20e-16, Fisher’s Exact test, Fig. 2E). Lamin B LAD-associated probes, while overrepresented as well, were less enriched (1.54-fold, P<2.20e-16, Fisher’s Exact test) and showed less methylation changes (Figs. 2E and Suppl. Fig. 3C). We also tested for the enrichment of transcription factor binding sites among the regions comprising the differentially methylated probes in HGPS using ELMER. In agreement with our ATAC-seq findings, AP1 family members were the most enriched in these regions (Odds ratio > 2.1, 95% confidence interval, Suppl. Fig. 3D), indicating that the transcription factor family is indeed at the center of epigenetic changes in HGPS fibroblasts.

Zhou et al. recently identified a subset of lamina-associated CpGs in PMDs (termed ‘solo-WCGWs’), which track the mitotic history of a cell and gain substantial age-related hypomethylation in a wide range of tissues^32^. Given the occurrence of different aging phenotypes in HGPS cells, we wondered whether these ‘solo-WCGWs’ CpGs show similar hypomethylation in HGPS. Intriguingly, median methylation levels of PMD-associated ‘solo-WCGWs’ were not reduced, but increased significantly in HGPS compared to control cells (P<2.20e-16, Welch Two Sample t-test, Fig. 2D). In comparison, methylation of ‘solo-WCGW probes associated with highly methylated domains (HMDs) exhibited only minor changes (Fig. 2D). Altogether, these results indicate that DNA methylation changes in HGPS cells occur primarily in lamina-associated, partially methylated regions and are distinct from changes found during normal aging.

### Epigenetic deregulation of LADs contributes to aberrant gene expression in HGPS

To investigate whether the identified LAD-specific chromatin accessibility and DNA methylation changes contribute to aberrant gene expression in HGPS cells, we performed RNA sequencing (RNA-seq) using fibroblasts from six HGPS patients and three controls. A total of 343 genes were found to be significantly (q<0.05, Benjamini-Hochberg) differentially expressed between HGPS and control cells, of which 160 were upregulated and 183 were downregulated in HGPS, respectively (Fig. 3A). Importantly, many of these genes, especially those more strongly deregulated, overlapped with genes previously identified to be differentially expressed in HGPS^11,41^ (Suppl. Fig. 4A).

**Figure 3.**
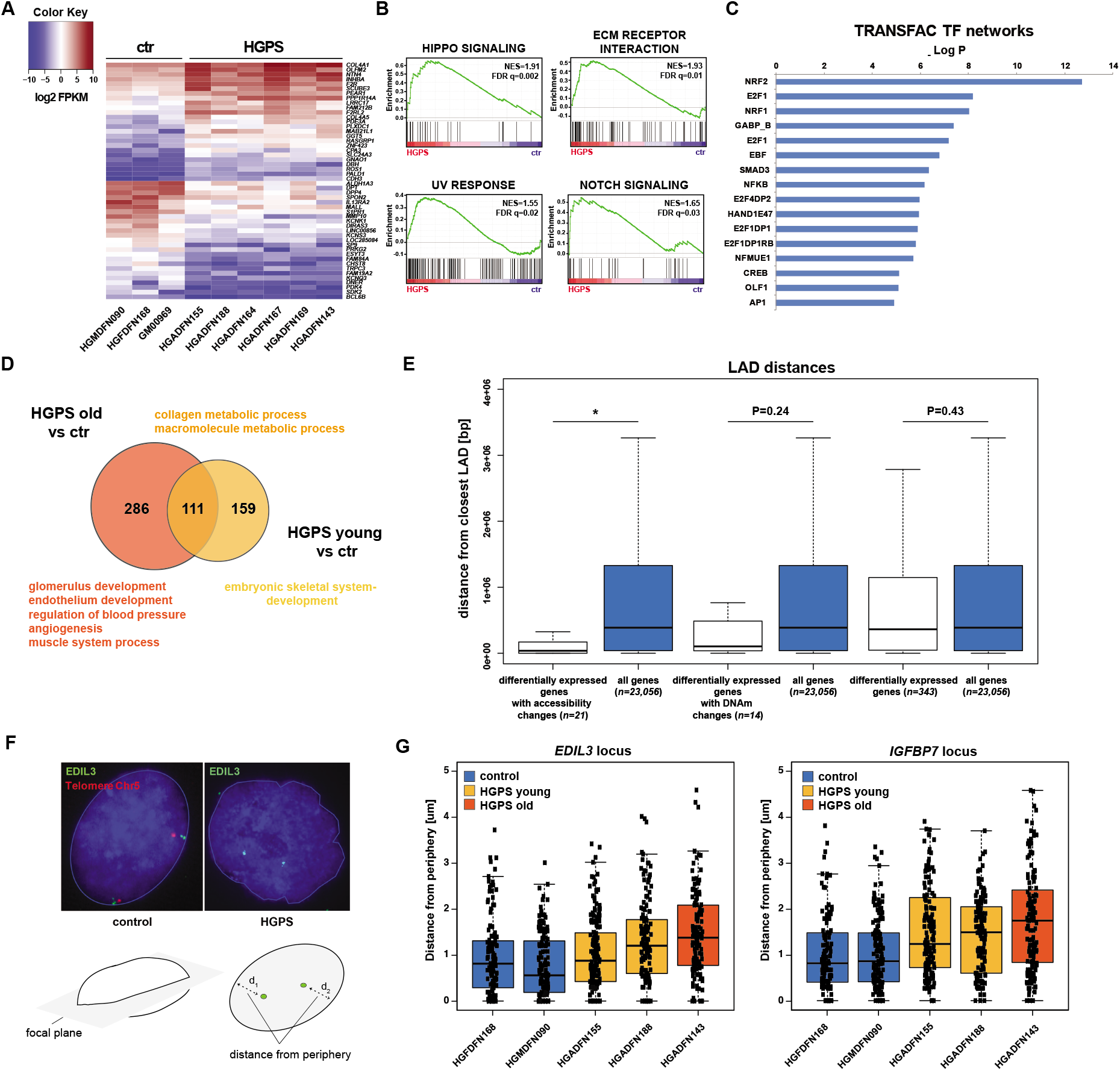
Epigenetic deregulation of lamina-associated domains (LADs) contributes to aberrant gene expression in HGPS. **(A)** 50 most differentially (q<0.05, Benjamini-Hochberg) expressed genes in six HGPS vs three control samples. Lowly expressed genes are shown in blue, highly expressed ones in red. FPKM = Fragments Per Kilobase of transcript per Million mapped reads. **(B)** Selection of Gene Ontology (GO), Kyoto Encyclopedia of Genes and Genomes (KEGG) and hallmark gene sets enriched (FDR q<0.05) in HGPS fibroblasts. NES=Normalized Enrichment Score. **(C)** TRANSFAC transcription factor (TF) network analysis of upstream factors controlling the observed expression changes. **(D)** Venn diagram showing numbers of genes overlapping between HGPS young (<8 years) vs. control samples and HGPS old (>8 years) vs control samples, respectively. The GO processes characteristic of each comparison are given. **(E)** Median distance to the nearest LAD (Lamin A and/or Lamin B) for the indicated sets of genes. Numbers of genes in each group and Wilcoxon rank sum test P-values (with continuity correction) are given. *P=2.25e-4. **(F)** Representative *EDIL3* FISH images in HGPS and control nuclei. A telomeric probe (red) on Chr5 was used as a positive staining control. The distance from the FISH signal to the nuclear periphery was measured in the focal plane in cells exhibiting a clear biallelic signal. **(G)** Quantification of (F) for *EDIL3* and *IGFBP7* loci in two control and three HGPS cell lines for 60 cells per sample. *EDIL3:* P=3.03e-08, Welch Two Sample t-test. *IGFBP7*: P=2.75e-13, Welch Two Sample t-test.

Gene Ontology (GO) analysis suggested ‘organismal’ and ‘developmental’ processes as well as ‘signaling’ and ‘cell communication’ as the main pathways to be affected by these changes (Suppl. Fig. 4B). In addition, Gene Set Enrichment Analyses (GSEA) highlighted Hippo signaling, ECM receptor interaction, UV response and Notch signaling, among others, to be significantly overrepresented in the set of differentially regulated genes (Fig. 3B and Suppl. Fig. 4C). TRANSFAC analysis of putative upstream regulators identified NRF2 as the main transcription factor associated with these changes (Fig. 3C). Importantly, NRF2 has previously been reported to be sequestered to the nuclear lamina by Progerin and to cause a subset of HGPS-associated expression changes^11^.

Given the rapid progression of age-related pathologies in HGPS patients, we also wondered whether cells from young and older patients differ significantly with regard to their expression patterns. Indeed, deregulated genes in younger patients (<8 years) were associated with the GO terms ‘embryonic skeletal system development’ and ‘anterior/posterior pattern specification’ (Fig. 3D and Suppl. Fig. 4D). In comparison, differentially expressed genes in older patients (>8 years) were associated with typical HGPS-related pathological features including ‘glomerulus development’, ‘endothelium development’, ‘regulation of blood pressure’, ‘angiogenesis’, and ‘muscle system process’ (Fig. 3D and Suppl. Fig. 4D). In conclusion, both HGPS-related developmental aberrancies and the worsening of disease pathology can be tracked at the level of gene expression *in vitro.*

To test whether LAD-associated epigenetic changes contribute to the gene expression patterns observed in HGPS fibroblasts, we compared the RNA-seq datasets with the DNA methylation and ATAC-seq datasets. Of the 343 genes with significant expression changes, 21 showed simultaneous changes in accessibility in HGPS (Suppl. Fig. 4E). In comparison, simultaneous DNA methylation changes were observed in 14 genes (Suppl. Fig. 4F), and three of the differentially expressed genes (*EDIL3, RELN* and *ZNF423*) were both differentially methylated and differentially accessible. Interestingly, the 21 genes undergoing both accessibility and expression changes are localized in or in close proximity to LADs in the genome of dermal fibroblasts (P=2.25e-4, Wilcoxon rank sum test, Fig. 3E). Although not reaching statistical significance, a similar trend was observed for the 14 genes undergoing both DNA methylation and expression changes (P=2.43e-1, Wilcoxon rank sum test, Fig. 3E). Taken together, these findings reveal an important subset of differentially expressed genes, which is affected by the epigenetic deregulation of LADs in HGPS.

To further confirm the epigenetic deregulation of LADs in HGPS, we performed fluorescence in situ hybridization (FISH) experiments in HGPS fibroblast nuclei. More specifically, we measured the distance of *EDIL3* and IGFBP7-specific FISH signals to the nuclear lamina (Fig. 3F, lower panel). The former encodes an integrin ligand with an important role in angiogenesis, as well as in vessel wall remodeling and development^42,43^; the latter encodes a member of the insulin-like growth factor-binding protein (IGFBP) family and is related to cellular senescence and modulation of angiogenesis^44–46^. Both *EDIL3* (P=3.03e-08, Welch Two Sample t-test) and *IGFBP7* (P=2.75e-13, Welch Two Sample t-test) were consistently localized farther away from the nuclear periphery in HGPS cells than in control cells (Figs. 3F and 3G), and we subsequently verified the increased expression of both genes in HGPS cells using quantitative RT-PCR (Suppl. Fig. 4G). These findings further support the notion that intranuclear re-localization of lamina-associated loci underpins the aberrant gene expression patterns in HGPS.

## Discussion

Previous studies have reported widespread histone modification changes in HGPS cells^15,18,19^, but little is known about their effect on chromatin accessibility and their relationship to DNA methylation patterns. To explore the nature and extent of potential alterations in these regulatory layers, as well as their influence on disease-specific gene expression, we subjected primary dermal fibroblasts from different HGPS patients to an integrated analysis using ATAC-see/-seq, DNA methylation profiling and RNA-seq. Our data reveal epigenetic deregulation of LADs as a novel, defining feature of the HGPS epigenome, which simultaneously contributes to aberrant gene expression in the disease.

We found HGPS-specific chromatin accessibility and DNA methylation changes to be significantly enriched in genomic regions that are in contact with the nuclear lamina in normal dermal fibroblasts^26,47^. This enrichment was observed for both regions gaining and losing accessibility in the disease, thus indicating considerable re-configuration of peripheral chromatin in HGPS nuclei. Importantly, as more than two thirds of the differentially accessible regions exhibited increased accessibility in the disease, lamina-associated chromatin appears to undergo substantial relaxation, which is in agreement with the observation that Progerin-expressing cells are characterized by reduced levels of the heterochromatin marks H3K9me3 and H3K27me3 in the vicinity of the nuclear lamina^18,19^. Similarly, Hi-C experiments with HGPS fibroblasts have demonstrated a considerable loss of chromatin compartmentalization, i.e., a reduction of chromatin compartment identity, at later passages^19,21^. Our finding that nuclear malformation in HGPS correlates with the loss of highly accessible chromatin in the nuclei of HGPS fibroblasts is in accordance with these observations.

The rather limited number of regions exhibiting significant chromatin accessibility changes may appear surprising, given the numerous epigenetic aberrations observed in HGPS cells. However, we emphasize that in Lamin A/C immunostainings HGPS fibroblasts were characterized by high population heterogeneity, with individual nuclei exhibiting a wide range of nuclear malformations. The set of regions we identified as significantly differentially accessible may therefore represent a consensus of chromatin regions shared by a minimum number of Progerin-expressing cells in the population. Future experiments with single-cell resolution should yield a better overview of the nature of these changes in individual cells.

LADs largely overlap with PMDs, which have been shown to exhibit reduced levels of DNA methylation as a consequence of mitotic aging^32^. Surprisingly, however, in HGPS cells we observed an increase of median methylation levels in these regions, specifically those associated with Lamin A. As most of the DNA methylation changes in HGPS cells occurred in gene bodies and intergenic regions (Fig. 2B), we interpret our results to reflect a HGPS-specific relocalization of formerly LAD-associated regions within the nucleus, leading to higher gene expression and concomitant DNA methylation changes as observed in the case of the *EDIL3* and *IGFBP7* loci. Such a conclusion is supported by the finding that gene expression positively correlates with increased gene body methylation^48^.

Our results also indicate that considerable DNA methylation differences exist not only between HGPS and control samples, but also within the group of patient fibroblasts. Intriguingly, these are manifested in a DNA methylation age acceleration for some patients, but not others (Fig. 2C). Despite small deviations, which are likely to be attributable to differences in the number of population doublings, our DNA methylation age estimates closely resemble recently published data for the same samples with the exception of HGADFN143^25^. However, we detected no or very limited differences between the two groups with regard to chromatin accessibility and gene expression, respectively, and the extent of these changes did not match those observed between HGPS and control cells (data not shown). While this is likely to be a consequence of the relatively low number of samples from both groups used for the comparisons, further studies are necessary to better decipher the molecular implications of the detected DNA methylation alterations in HGPS cells.

Although HGPS fibroblasts exhibited a substantial number of differentially expressed genes, the effect of LAD-related chromatin accessibility and DNA methylation changes on gene expression was limited. While this is in line with the observation that LADs are generally gene-poor^26^, it also suggests that other mechanisms are likely to contribute to the aberrant gene expression of HGPS cells. In this regard, NRF2 mislocalization to the nuclear lamina has recently been demonstrated to be a key contributor to the HGPS-specific transcriptome^11^. Our data are in agreement with this finding, since we identified the transcription factor as one of the key factors responsible for the HGPS-related gene expression changes, even though its target genes did not reach statistical significance (FDR q-val<0.05) in our GSEA analyses (Suppl. Fig. 4C). Furthermore, NRF2 binding motifs were significantly (q=1.80e-3, Benjamini-Hochberg) enriched in the differentially accessible regions between HGPS and control cells (Suppl. Fig. 2E). Because the transcription factor has been shown to be sequestered to the nuclear lamina by Progerin^11^, it is tempting to speculate that the epigenetic changes observed in LADs are involved in the NRF2-mediated differential expression in the disease.

The nuclear lamina is known to be a resting place for transcription factors, serving to sequester them away from chromatin^49^. c-Fos, a member of the AP1 family of transcription factors, for example, is negatively regulated by binding to Lamin A/C at the nuclear lamina^36^. This function might be impaired by the presence of Progerin in HGPS cells, since we found AP1 members to constitute the transcription factor family that is most affected by the epigenetic deregulation in HGPS (Fig. 1G, Suppl. Fig. 3D). Intriguingly, both *IGFBP7* and *EDIL3* contain consensus binding sites for AP1 family members in their respective promoter or enhancer^45,50^, further indicating that their differential expression is likely driven by epigenetic changes in HGPS cells.

In summary, our results suggest a scenario, in which the Progerin-driven nuclear malformation of HGPS nuclei causes substantial, but potentially locally stochastic epigenetic re-configuration of LAD-specific chromatin (Fig. 4). As peripheral heterochromatin is diminished in these cells^15,18,19^, many of the affected regions gain a more relaxed chromatin environment that is more permissive to the binding of transcription factors and might thus facilitate differential expression. In some cases, chromatin decondensation and disease-specific differential expression of formerly LAD-associated loci accompanies their relocalization within the nucleus (Fig. 4), as detected for *EDIL3* and *IGFBP7.* Further experiments are needed to clarify whether similar mechanisms contribute to disease-specific gene expression in other tissues and whether they are also active in classical aging.

**Figure 4.**
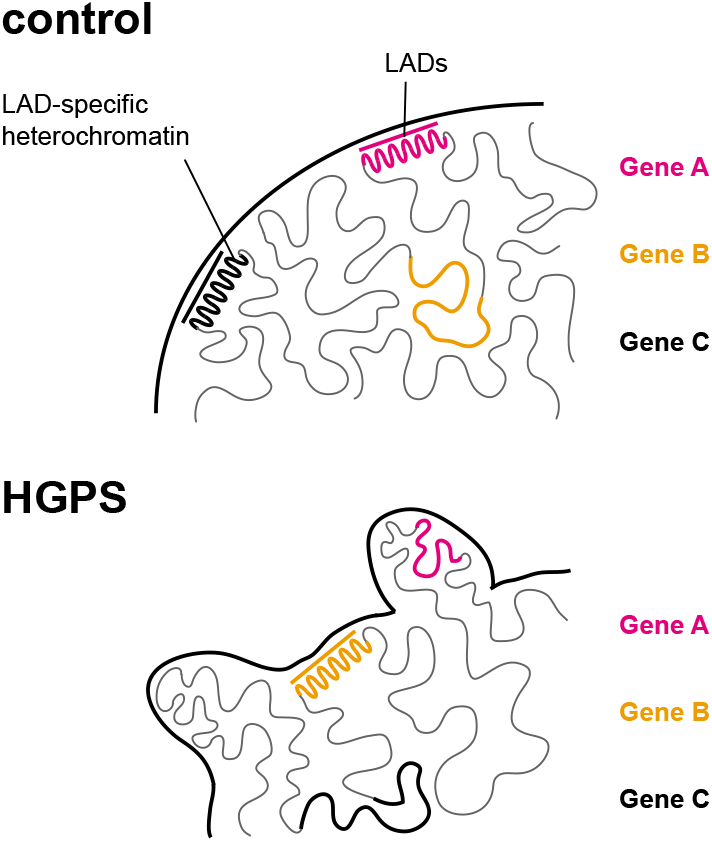
Epigenetic deregulation of lamina-associated domains (LADs) in HGPS. Progerin-driven nuclear malformation in HGPS nuclei causes substantial, but potentially locally stochastic epigenetic reconfiguration of LAD-specific chromatin. As peripheral heterochromatin is diminished in these cells, many of the affected regions gain a more relaxed chromatin environment that is more permissive to the binding of transcription factors and might thus facilitate differential expression. In some cases, chromatin decondensation and disease-specific differential expression of formerly LAD-associated loci coincides with their relocalization within the nucleus, as detected in the case of *EDIL3* and *IGFBP7.*

## Methods

### Samples

Primary HGPS patient (HGADFN155, HGADFN271, HGADFN188, HGADFN164, HGADFN122, HGADFN178, HGADFN167, HGADFN169 and HGADFN143) and parental skin fibroblasts (HGMDFN090 and HGFDFN168) were obtained from the Progeria Research Foundation (Boston, MA, USA). Age-matched primary skin fibroblasts (GM05659, GM02036, GM01864 and GM00969) were obtained from the Coriell Cell Repository (Camden, NJ, USA). A detailed overview of the cells used for each experiment is given in Suppl. Tab. 1. Cells were grown in DMEM high glucose medium supplemented with 10% fetal bovine serum (FBS) and 1% penicillin/streptomycin under standard 37°C and 5% CO_2_ conditions.

### ATAC-see

Hyperactive Tn5 transposase production and Tn5 transposome assembly using Atto-590N-labeled oligonucleotides were carried out as described previously^33^. Cells were grown on cover slips until 70-80% confluence and fixed with 3.8% paraformaldehyde (PFA) for 10 minutes at room temperature. They were then permeabilized with lysis buffer (10 mM TRIS-HCl pH 7.4, 10 mM NaCl, 3 mM MgCl_2_, 0.01% IGEPAL CA-630), washed with 1x phosphate buffered saline (PBS) twice and incubated with the transposome mixture (100 nM assembled Tn5-Atto-590N-transposomes, 25 μl Nextera tagmentation buffer, ddH_2_O to 50 μl) for 30 minutes at 37°C. Subsequently, cover slips were washed three times (15 minutes each) with 1x PBS containing 0.01% SDS and 50 mM EDTA at 55°C and immunostained. Imaging was performed using an Olympus FluoView FV1000 microscope. For each sample, 50 cells from three technical replicates were analyzed to determine nuclear malformation and the presence of ATAC-see foci. A nucleus was scored as malformed when showing lobulation characteristic of HGPS cells.

### ATAC-seq

ATAC-seq was performed as described previously^51^. Briefly, 50,000 cells were washed with ice cold 1x PBS and resuspended in 50 μl lysis buffer (10 mM TRIS-HCl pH 7.4, 10 mM NaCl, 3 mM MgCl_2_, 0.1% IGEPAL CA-630). The lysis reaction was carried out while spinning down the samples at 500 g for 10 minutes at 4°C. Samples were then resuspended in transposition buffer (25 μl 2x TD buffer (Illumina), 2.5 μl TDEI (Tagment DNA Enzyme, Illumina) and 22.5 μl ddH_2_O) and incubated for 30 minutes at 37°C. Subsequently, samples were purified using the MinElute PCR Purification kit (Qiagen). Final libraries were then PCR-amplified, purified once again with the same kit and subjected to paired-end sequencing on a HiSeq 4000 platform (Illumina).

Reads were trimmed by removing stretches of bases having a quality score of <30 at the ends of the reads. The reads were mapped using Bowtie 2^52^ against the hg19 assembly of the human genome. Peaks were called using MACS2^53^ and differential peaks were quantified by DESeq2^54^.

Distribution of significant, non-sex chromosome-associated ATAC-seq peaks across the genome was determined using the subsetByOverlaps and Genomic Regions functions with the TxDb.Hsapiens.UCSC.hg19.knownGene Bioconductor annotation package (version 3.2.2) in R (version 3.3.1). The overlap with histone modifications and Lamin A and Lamin B LADs, respectively, was calculated in the same way using previously published ENCODE (ENCSR000ARX, ENCSR000ARV, ENCSR000APR, ENCSR000APN, ENCSR000APP, ENCSR000APQ, ENCSR000APO) and ChIP-seq data^26,47^. Poised enhancers for dermal fibroblasts were defined as regions containing pairs of H3K4me1 peaks in close proximity (<1,500 bp) using ENCODE ChIP-seq data (ENCSR000ARV); active enhancers were obtained directly from ENCODE (ENCSR871EJM). Transcription factor binding sites overlapping with significant, non-sex chromosome-associated ATAC-seq peaks were determined using the HOMER motif analysis tool^55^.

### DNA methylation analysis

DNA methylation profiles were generated using Infinium MethylationEPIC BeadChips (Illumina), following the manufacturer’s instructions. Methylation data analysis was carried out using the R Bioconductor package Minfi (v1.20.2)^56^. Specifically, raw .IDAT files were read and preprocessed. Methylation loci (probes) were filtered for high detection p-value (P>0.01, as provided by Minfi), location on sex chromosomes, ability to self-hybridize, and potential SNP contamination. Array normalization was performed using the *preprocessFunnorm* function, available in Minfi^56^. Quality control was performed after every preprocessing step. Subsequently, differentially methylated probes were identified by fitting a linear model followed by statistical analysis using an empirical Bayes method to moderate standard errors. Lastly, differentially methylated probes were filtered by significance threshold (P<0.05, F-test, after correction for multiple testing using the Benjamini-Hochberg method).

For consensus clustering, probe clusters were identified using the Minfi function boundedClusterMaker with a maximum cluster width of 1,500 bp and a maximum gap of 500 bp. Using the ConsensusClusterPlus package^57^, consensus clustering was performed with the 5,000 most variable probe clusters, i.e., the 5,000 probe clusters having the highest standard deviation, with the following parameters: maxK = 6; reps = 1,000; pItem = 0.8; and pFeature = 1. Samples were then assigned to the optimal number of clusters, and β values were sorted by hierarchical clustering for visualization. Finally, a Principle Component Analysis (PCA) was generated on the basis of the identified 5,000 probe clusters with the R package FactoMineR^58^.

For the comparison of LAD-and solo-WCGW CpG probe methylation levels between HGPS and control samples, we used previously published locations of Lamin A LADs^47^, Lamin B LADs^26^ and solo-WCGWs CpGs^32^. The enrichment of transcription factor binding sites among the differentially methylated regions was analyzed using ELMER 2.0^59,60^ with a minimum motif quality ‘B’ and a minimum incidence of 10. Finally, DNA methylation age estimates were obtained using a recently published algorithm (Skin&Blood Clock^25^) and corrected for passage number using a passage factor ρ, with ρ = passage number*(3.32*log(cells harvested/cells seeded)).

### DNA FISH

Bacterial Artificial Chromosomes (BACs) were obtained from the Children’s Hospital Oakland Research Institute (CHORI) (Oakland, CA, USA). Suppl. Tab. 2 contains a list with all BACs used and the genes they cover. BACs were labeled with the ENZO Nick translation DNA labeling system 2.0 (ENZO) using SEEBRIGHT Red 580 dUTP (ENZO) and SEEBRIGHT Green 496 dUTP (ENZO), following the manufacturer’s instructions. Subsequently, 500 ng of each probe were precipitated with 5 μl of Cot-1 DNA (1 mg/ml) (Invitrogen) and resuspended in 15 μl of hybridization buffer (10% dextran sulfate, 50% formamide, 2x SSC pH 7). Ultimately, FISH was performed as previously described^61^. Slides were imaged using Zeiss Axioskop 2 and Olympus FluoView FV1000 microscopes. For each probe, images were acquired for 60 cells. Distance measurements were performed with Fiji^62,63^ using confocal sections with a clear FISH signal. For group-specific comparisons, the respective HGPS and control sample data were pooled and subjected to a Welch Two Sample t-test with a 95% confidence interval in R.

### Immunostainings

For Lamin A immunostainings, cells were grown on coverslips, fixed with 3.8% PFA in 1x PBS for 10 minutes at room temperature and permeabilized with 0.3% Triton X-100 in 1x PBS for 15 minutes at room temperature. Coverslips were subsequently washed three times with 1x PBS (10 minutes each) and blocked with 10% FBS in 1x PBS with 0. 1% (v/v) Tween 20 (PBT) for 1 h at room temperature. They were then incubated with a 1:250 dilution of Lamin A/C primary antibody (sc7292, Santa Cruz) in the same solution for 90 minutes, washed three times with 1x PBS (10 minutes each) and incubated with a 1:500 dilution of Alexa 488 secondary antibody (A11017, ThermoFisher) in the same solution for 45 minutes at room temperature. Finally, coverslips were washed three times with 1x PBS (10 minutes each), stained with DAPI and mounted onto slides. Slides were imaged using Zeiss Axioskop 2 and Olympus FluoView FV1000 microscopes. For quantification of malformed nuclei, severely misshapen nuclei (on the basis of significant blebbing and wrinkling) were determined in three technical replicates with 100 cells counted per replicate.

### RNA sequencing

RNA was isolated using TRIzol (Invitrogen) and following the manufacturer’s instructions. Total RNA was then purified with the RNA Clean & Concentrator-5 kit (Zymo Research) and reverse-transcribed using SuperScript III reverse transcriptase (Invitrogen), again following the manufacturer’s instructions. Libraries were prepared with the TruSeq RNA sample preparation kit (Illumina) and sequenced on a HiSeq 4000 machine (Illumina) using 50 bp single reads.

Reads were trimmed by removing stretches of bases having a quality score of <30 at the ends of the reads. The reads were mapped using Tophat 2.0.6^64^ against the hg19 assembly of the human genome. Differential expression was quantified using DESeq2^54^ and Cuffdiff 2.0^65^ and subjected to multiple testing corrections. Genes with a q-value smaller than 0.05 were considered differentially expressed.

Gene ontology (GO) analyses were performed using the AmiGO 2 database^66–68^. TRANSFAC analyses were carried out using the TRANSFAC® Public 6.0 database in Match – 1.0^69^. Gene Set Enrichment Analysis (GSEA)^70^ was performed utilizing the RNA sequencing datasets of HGPS and control fibroblasts and the ‘hallmark’ (v5.0, Arthur Liberzon^71^, Broad Institute) and ‘KEGG’ (KEGG (Kyoto Encyclopedia of Genes and Genomes)) collections from the Molecular Signature Databases (MSigDB) supplemented with the ‘NRF2_01’ (v6.0, Xiaohui Xie, Broad Institute) gene set, using the parameters ‘gene_set permutation’ and ‘1000 permutations’. Gene signatures with a false discovery rate (FDR) q-value of < 0.05 were considered enriched.

## Acknowledgements

We thank the DKFZ Genomics and Proteomics Core Facility for sequencing and methylation services. We also thank Leslie Gordon, Susan Campbell, Wendy Norris and the Progeria Research Foundation for generously providing HGPS cells and information. This work was supported by a research grant from the Helmholtz program ‘Aging and Metabolic Programming’ (AMPro) to F.L..

## Author Contributions

F.K., F.B., G.R., J.G., F.L. and M.R.-P. analyzed the data. F.K. performed the experiments assisted by T.M. F.L. and M.R.-P. conceived the study. F.K., F.L. and M.R.-P. wrote the manuscript. All authors read and approved the submitted version.

