## Supplementary material for "Epigenetic deregulation of lamina-associated domains in Hutchinson-Gilford Progeria Syndrome" (from another version of this preprint: version 2, 2019-01-21)

**Supplementary Table 1:** Primary cells used in this study

PRF = Progeria Research Foundation, CCR = Coriell Cell Repository, 1 = DNA methylation profiling, 2 = ATAC-see, 3 = ATAC-seq, 4 = RNAseq

| Sample ID | Status | Gender | Age [years] | Source | Experiments |
|---|---|---|---|---|---|
| HGADFN155 | HGPS | Female | 1.17 | PRF | 1,2,3,4 |
| HGADFN271 | HGPS | Male | 1.25 | PRF | 1,3 |
| HGADFN188 | HGPS | Female | 2.25 | PRF | 1,2,3,4 |
| HGADFN164 | HGPS | Female | 4.67 | PRF | 1,4 |
| HGADFN122 | HGPS | Female | 5.00 | PRF | 1 |
| HGADFN178 | HGPS | Female | 6.92 | PRF | 1 |
| HGADFN167 | HGPS | Male | 8.42 | PRF | 1,3,4 |
| HGADFN169 | HGPS | Male | 8.50 | PRF | 1,3,4 |
| HGADFN143 | HGPS | Male | 8.83 | PRF | 1,3,4 |
| GM05659 | control | Male | 1.00 | CCR | 1,2 |
| GM00969 | control | Female | 2.00 | CCR | 1,4 |
| GM01864 | control | Male | 11.00 | CCR | 1 |
| GM02036 | control | Female | 11.00 | CCR | 1 |
| HGMDFN090 | control | Female | 37.83 | PRF | 1,3,4 |
| HGFDFN168 | control | Male | 40.42 | PRF | 1,2,3,4 |

**Supplementary Table 2:** BACs used in this study

| BAC Name | Reference | Chromosome | Location |
|---|---|---|---|
| EDIL3 | RP11-845G7 | 5 | *EDIL3* gene |
| Chr5 control | RP11-82M24 | 5 | telomere |
| IGFBP7 | RP11-589G9 | 4 | *IGFBP7* gene |
| RELN | RP11-57M15 | 7 | *RELN* gene |
| SOX11 | RP11-103F8 | 2 | *SOX11* gene |
| SMAD9 | RP11-354L15 | 13 | *SMAD9* gene |
| KCNK1 | RP11-349N15 | 1 | *KCNK1* gene |

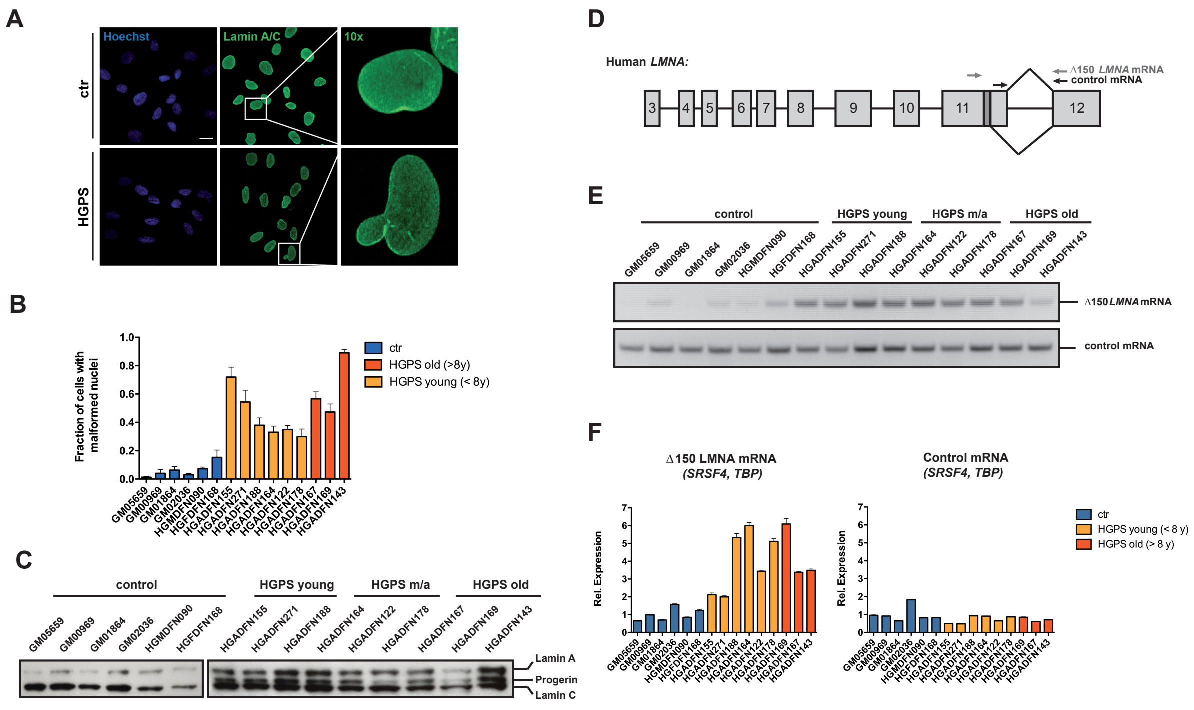


**Suppl. Figure 1. Nuclear morphology and Progerin expression in fibroblast model system. (A)** α-Lamin A/C immunofluorescence showing characteristic nuclear malformation in HGPS (HGPS: HGADFN188, ctr: HGMDFN090, scale bar = 10 µm). **(B)** Quantification of malformed nuclei in HGPS and control fibroblast cell lines. HGPS cells were grouped as young (<8 years) or old (>8 years) depending on the age of the donor (source: Progeria Research Foundation). **(C)** Western Blot with α-Lamin A/C antibody (sc7292, Santa Cruz) showing expression of Progerin in HGPS samples. The band corresponding to Lamin C was used as a loading control. **(D)** Schematic representation of *LMNA* gene. The locations of the primers designed for the detection of the Δ150 *LMNA* and control mRNAs are indicated. **(E)** Detection of Δ150 *LMNA* mRNA in HGPS fibroblasts through RT-PCR (50 cycles). **(F)** Expression of Δ150 *LMNA* mRNA relative to SRSF4 and TBP in HGPS and control fibroblasts as measured through RT-qPCR.

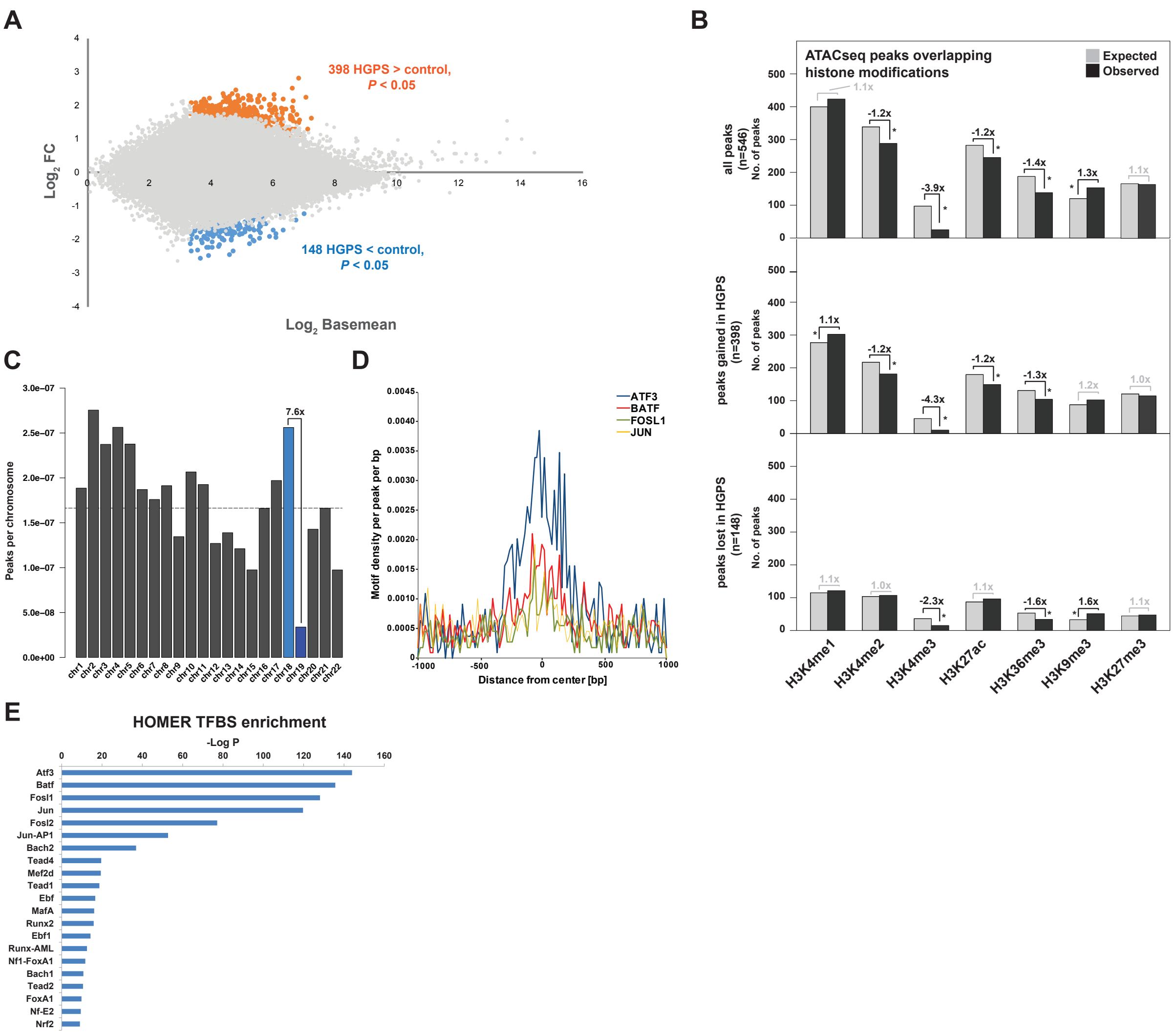


**Suppl. Figure 2. ATAC-see and ATAC-seq reveal genome-wide chromatin accessibility changes enriched in lamina-associated domains (LADs). (A)** Log2 Fold Change (FC) and log2 basemean of regions gaining (n=398) and losing (n=148) accessibility in HGPS, respectively (q<0.05, Benjamini-Hochberg). **(B)** Overlap of ATAC-seq peaks with the indicated histone modifications (*P<0.05, Fisher's Exact test). The expected number of peaks was calculated based on the fraction of all (including non-significant (q<0.05, Benjamini-Hochberg)) peaks overlapping with a certain histone modification normalized to the number of peaks significantly (q<0.05, Benjamini-Hochberg) gaining or losing accessibility, or both. **(C)** Distribution of ATAC-seq peaks across chromosomes (normalized to chromosome length) with mean number indicated as a dashed line (mean=1.66e-07 peaks per chromosome). Chromosome 18 exhibited 7.6 times more differentially accessible peaks than chromosome 19. **(D)** Motif density plot for selected transcription factors. **(E)** NRF2 binding motifs are among the transcription factor binding sites enriched in the differentially accessible regions (q=1.80e-3, Benjamini-Hochberg).

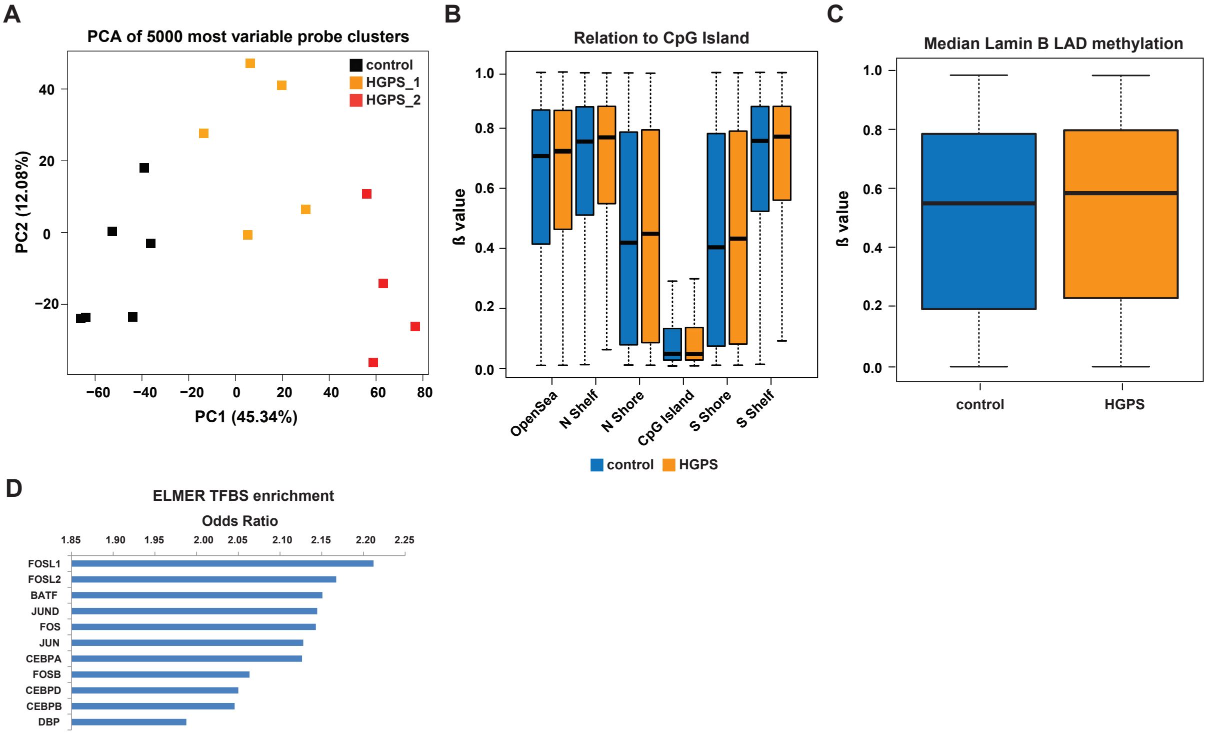


**Suppl. Figure 3. DNA methylation profiling in HGPS reveals two patient subgroups and lamina-associated domain (LAD)-enriched hypermethylation. (A)** Principal component analysis (PCA) of 9 HGPS and 6 control samples using the 5000 most variable probe clusters. The two HGPS subgroups identified through this analysis are indicated as HGPS_1 and HGPS_2. **(B)** Median methylation (β value) levels of OpenSea (P=1.49e−93), Shelf (N: P=3.17e−09, S: P=8.77e−08), Shore (N: P=2.5e−08, S: P=1.17e−07) and CpG Island (P=0.82)-associated probes in control (blue) and HGPS (orange) samples (all: F test, Benjamini-Hochberg). **(C)** Median methylation (β value) levels of Lamin B LAD-associated probes in control (blue) and HGPS (orange) samples (P=2.20e-16, Welch Two Sample t-test). **(D)** ELMER Transcription Factor Binding Site (TFBS) enrichment analysis reveals that members of the AP1 family are enriched in the differentially methylated regions (95% Confidence Interval).

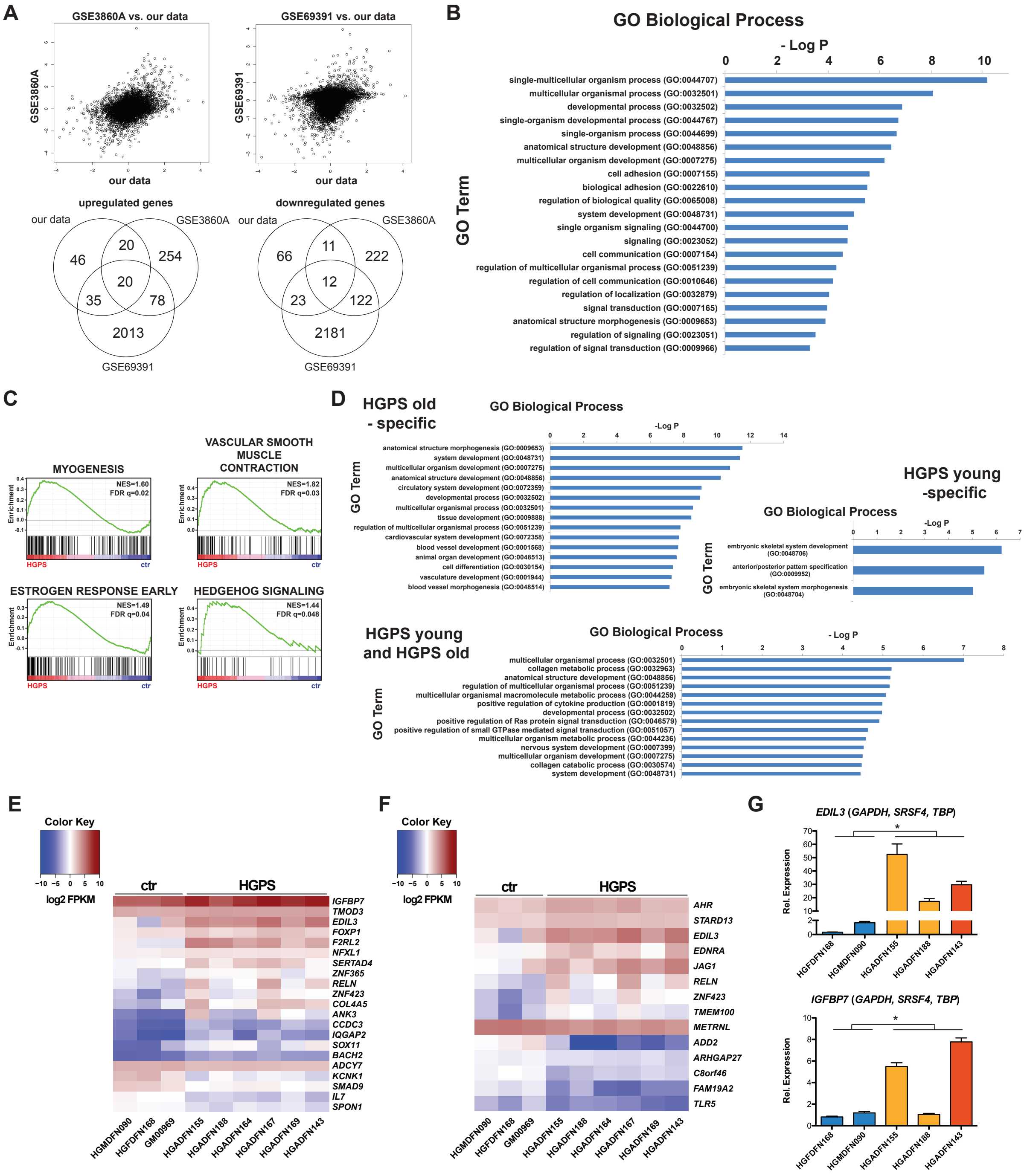


**Suppl. Figure 4. Epigenetic deregulation of lamina-associated domains (LADs) contributes to aberrant gene expression in HGPS. (A)** Comparison of RNAseq data with previous HGPS expression studies. Log Fold Changes (FC) of genes in GSE3860A vs. our data (Pearson correlation r=0.37, P<2.20e-16) and in GSE69391 vs. our data (Pearson correlation r=0.23, P<2.20e-16), respectively are given in upper panels. Lower panels show the overlap of genes up- or downregulated, respectively, between our and the aforementioned studies. **(B)** Gene Ontology (GO) processes enriched among the differentially (q<0.05, Benjamini-Hochberg) expressed genes in HGPS. **(C)** Additional hallmark and Kyoto Encyclopedia of Genes and Genomes (KEGG) gene sets enriched (FDR q<0.05) in HGPS fibroblasts. **(D)** GO processes enriched among the differentially (q<0.05, Benjamini-Hochberg) expressed genes between HGPS old (>8 years) vs. control, HGPS young (<8 years) vs. control or HGPS old vs. young, respectively. **(E)** Differentially (q<0.05, Benjamini-Hochberg) expressed genes that show concomitant HGPS-specific chromatin accessibility changes in their promoter or gene body. Lowly expressed genes are shown in blue, highly expressed ones in red. FPKM = Fragments Per Kilobase of transcript per Million mapped reads. **(F)** Differentially ($q<0.05$, Benjamini-Hochberg) expressed genes that show concomitant HGPS-specific DNA methylation changes in their promoter or gene body. Lowly expressed genes are shown in blue, highly expressed ones in red. FPKM as in (E). **(G)** *EDIL3* and *IGFBP7* expression levels relative to those of *GAPDH*, *SRSF4* and *TFB* in control (blue), HGPS young (<8 years, orange) and HGPS old (>8 years, red) cells as measured in RT-qPCR (**EDIL3*: P=2.00e-3, *IGFBP7*: P=4.00e-4, unpaired t-test).
